# Turicibacterales protect mice from severe *Citrobacter rodentium* infection

**DOI:** 10.1101/2023.04.25.538270

**Authors:** Kristen L. Hoek, Kathleen G. McClanahan, Yvonne L. Latour, Nicolas Shealy, M. Blanca Piazuelo, Bruce A. Vallance, Mariana X. Byndloss, Keith T. Wilson, Danyvid Olivares-Villagómez

**Affiliations:** Department of Pathology, Microbiology, and Immunology, Vanderbilt University Medical Center, Nashville, TN, USA; Division of Gastroenterology, Hepatology, and Nutrition, Department of Medicine, Vanderbilt University Medical Center, Nashville, TN, USA; Center for Mucosal Inflammation and Cancer, Vanderbilt University Medical Center, Nashville, TN, USA; Department of Pediatrics, BC Children’s Hospital, University of British Columbia, Vancouver, BC, Canada; Vanderbilt Institute of Infection, Immunology, and Inflammation, Vanderbilt University Medical Center, Nashville, TN, USA; Vanderbilt Microbiome Innovation Center, Vanderbilt University Medical Center, Nashville, TN, USA; Veternas Affairs Tennessee Valley Healthcare System, Nashville, TN, USA; ORCID 0000-0002-1158-8976

## Abstract

One of the major contributors to child mortality in the world is diarrheal diseases, with an estimated 800,000 deaths per year. Many pathogens are causative agents of these illnesses, including the enteropathogenic (EPEC) or enterohemorrhagic (EHEC) forms of *Escherichia coli*. These bacteria are characterized by their ability to cause attaching and effacing lesions in the gut mucosa. Although much has been learned about the pathogenicity of these organisms and the immune response against them, the role of the intestinal microbiota during these infections is not well characterized. Infection of mice with *E. coli* requires pre-treatment with antibiotics in most mouse models, which hinders the study of the microbiota in an undisturbed environment. Using *Citrobacter rodentium* as a murine model for attaching and effacing bacteria, we show that C57BL/6 mice deficient in granzyme B expression are highly susceptible to severe disease caused by *C. rodentium* infection. Although a previous publication from our group shows that granzyme B-deficient CD4^+^ T cells are partially responsible for this phenotype, in this report we present data demonstrating that the microbiota, in particular members of the order Turicibacterales, have an important role in conferring resistance. Mice deficient in *Turicibacter sanguinis* have increased susceptibility to severe disease. However, when these mice are co-housed with resistant mice, or colonized with *T. sanguinis*, susceptibility to severe infection is reduced. These results clearly suggest a critical role for this commensal in the protection against entero-pathogens.

## INTRODUCTION

One of the major contributors to child mortality is diarrheagenic diseases caused by infectious pathogens, which cause approximately 800,000 deaths per year in children under the age of five (1, 2), with prevalent morbidity in adults (3). One such pathogen is *Escherichia coli*, in particular enteropathogenic (EPEC) or enterohemorrhagic (EHEC) forms. These bacteria utilize attaching and effacing (A/E) lesions to colonize the intestines of the host, leading to impaired ion and water transport across the intestinal epithelium (3, 4). Despite the clinical importance of these bacteria, their study in animal models is hindered by the requirement to treat the hosts with antibiotics prior to colonization (5), which disrupts the intestinal microbiota and its potential interactions with both the pathogen and intestinal immune cells.

*Citrobacter rodentium* is the only known naturally occurring A/E pathogen of mice, and it has become the principal mouse model for the study of A/E enteropathogenic diseases, providing invaluable knowledge about host-pathogen interaction mechanisms applicable to human EPEC and EHEC pathogens (6). Infection with *C. rodentium* usually results in transient colonization and inflammation of the distal colon, with little to no mortality in most mouse strains, such as C57BL/6, NIH Swiss, and BALB/c. A few strains, such as C3H/Hej, display high mortality, indicating that the genetic make-up of the host is critical for conferring protection against severe *C. rodentium* infection (4).

Many factors are involved in the host response against *C. rodentium* infection, including activation of the NOD2 sensor (7), production of IL-22 (8), IL-6 (9), and IL-23 (ref in (10)), innate immune response by ILC3 (key components of the initial response towards this pathogen, primarily by secreting IL-22 and promoting neutrophil recruitment (11)), and specific CD4^+^ T cell responses which become the main producers of IL-22 and IL17A (10). The microbiota is a critical component against *C. rodentium* infection. For example, it is well established that the intestinal mucus barrier is critical for protection against *C. rodentium* infection (12), and thus, microbial groups that promote the formation of this barrier are of great importance. Commensal organisms can also outcompete *C. rodentium* growth by utilizing key metabolites, such as monosaccharides (10). Although a few commensals have shown to be of critical importance against *C. rodentium*, there is still a significant gap identifying other beneficial microorganisms that may confer resistance against this pathogen.

*Turicibacter sanguinis* is the best studied species of the order Turicibacterales. This commensal anaerobe is prevalent in the intestinal microbiota of humans, mice, cows, pigs, and chickens (13–17). In mice intestines, *T. sanguinis* is fairly conspicuous, reaching relative frequencies above 20% (18), while in human fecal microbiota, *Turicibacter* frequencies are ∼0.5% (19). *Turicibacter* prevalence in mice and humans suggests an important role of this commensal in host physiology. Indeed, recent studies have indicated that the human *T. sanguinis* isolate MOL361 modifies the host lipidome, altering serum cholesterol and triglycerides in mice (20). Here, we present evidence that susceptibility to severe *C. rodentium* infection is associated with absence of Turicibaterales in the intestinal microbiota, and that protection from disease can be transferred by colonization with *Turicibacter sanguinis*.

## RESULTS

While studying the role of granzyme B, a potent serine protease primarily involved in cell-mediated cytotoxicity (21), we observed that infection of granzyme B-deficient (*Gzmb*^-/-^) mice with *Citrobacter rodentium* resulted in prominent weight loss, increased signs of disease (such as loose stool, diarrhea, rectal bleeding), scruffiness, and death (22), whereas control C57BL/6 wildtype (WT) mice showed little to no signs of disease with almost 100% survival. As we have previously reported, part of the phenotype observed in infected *Gzmb*^-/-^ mice is due to increased CD4^+^ T cell intrinsic pathogenicity (22). Most of the experiments in our previous report were performed with isolated mouse lines, i.e., *Gzmb*^-/-^ and WT mice were bred and kept independent from each other’s line. Because of the nature of the isolated mouse lines, it is possible that the severe disease phenotype observed was due in part to genetic drift or differences in the intestinal microbiota. To investigate this possibility, we crossed *Gzmb*^-/-^ with WT mice to generate *Gzmb*^+/-^ mice. F1 *Gzmb*^+/-^ mice were crossed among themselves to generate *Gzmb*^+/+^, *Gzmb*^+/-^, and *Gzmb*^-/-^ littermates. Susceptibility to severe *C. rodentium* infection was lost in littermate *Gzmb*^-/-^ mice, displaying similar weight and decreased disease signs to *Gzmb*^+/-^ or WT mice throughout the course of the experiment (**Figure 1A, B**). Colon injury was also reduced in littermate *Gzmb*^-/-^ mice in comparison to *Gzmb*^-/-^ mice from the isolated line (**Figure 1C**). Despite these differences, *C. rodentium* colonization was similar among these two groups (**Figure 1D** and data presented in our previous publication (22)). Thus, there are two *Gzmb*^-/-^ mouse lines based on their outcome to severe *C. rodentium* infection: susceptible (S, from the isolated line) and resistant (R, from littermates). Because weight loss closely corresponds to increased susceptibility, it will subsequently be used as the primary readout parameter.

**Figure 1.**
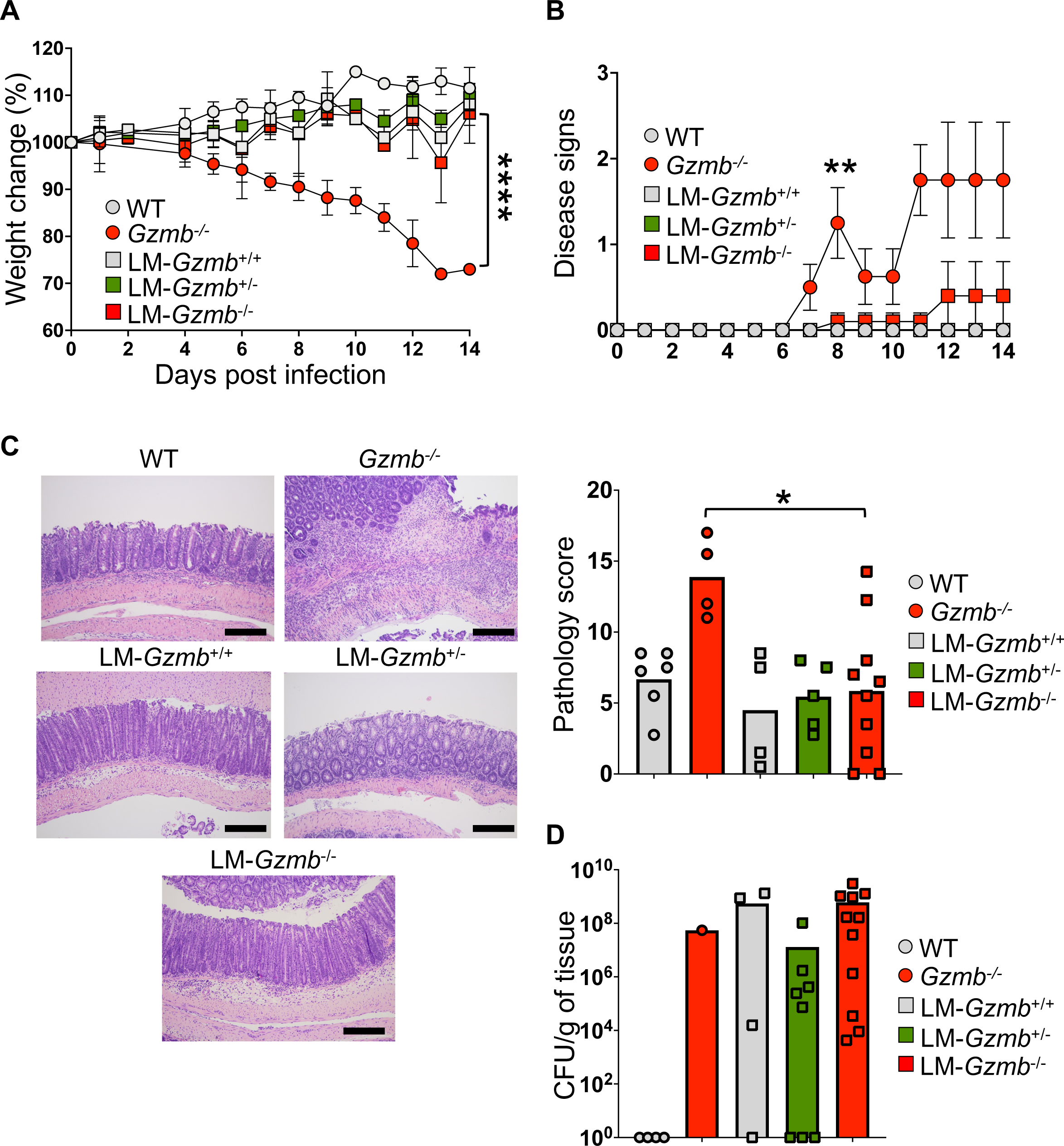
*Gzmb*^-/-^ littermate mice are resistant to severe *C. rodentium* infection. Littermate (LM) mice were infected with 5-10 × 10^8^ CFU, and monitored daily for weight (A), and signs of disease, such as diarrhea, scruffiness, and rectal irritation/bleeding (B). At day 14-15, colon injury (C) and *C. rodentium* colonization (D) were determined. n= 4-11. Data represent at least 3 independent experiments. For (A), ****P<0.001, Student’s t test for each time point after day 8. For (B) **<0.01, Student’s t test. For (C), *P<0.05, One-Way ANOVA. Bar indicates 250μm.

The differences in disease susceptibility in the two *Gzmb*^-/-^ mouse lines could be attributed to genetic drift or the intestinal microbiota. To discern between these two possibilities, (S)*Gzmb*^-/-^ mice were co-housed starting at weaning age with a single (R)*Gzmb*^-/-^ or (S)*Gzmb*^-/-^ mouse from a different litter. Mice are coprophagic, thus, co-housed animals are exposed to other mice’s fecal material, allowing for microbiota exchange. Susceptible mice cohoused with (R)*Gzmb*^-/-^ mice maintained their starting weight similar to WT mice (**Figure 2A**). In all experiments performed, susceptibility to severe disease was never transmitted to resistant mice, indicating that resistance is a dominant trait. Thus, the intestinal microbiota and not genetic drift, is the most likely factor responsible for the differences in disease susceptibility between the two *Gzmb*^-/-^ mouse lines.

**Figure 2.**
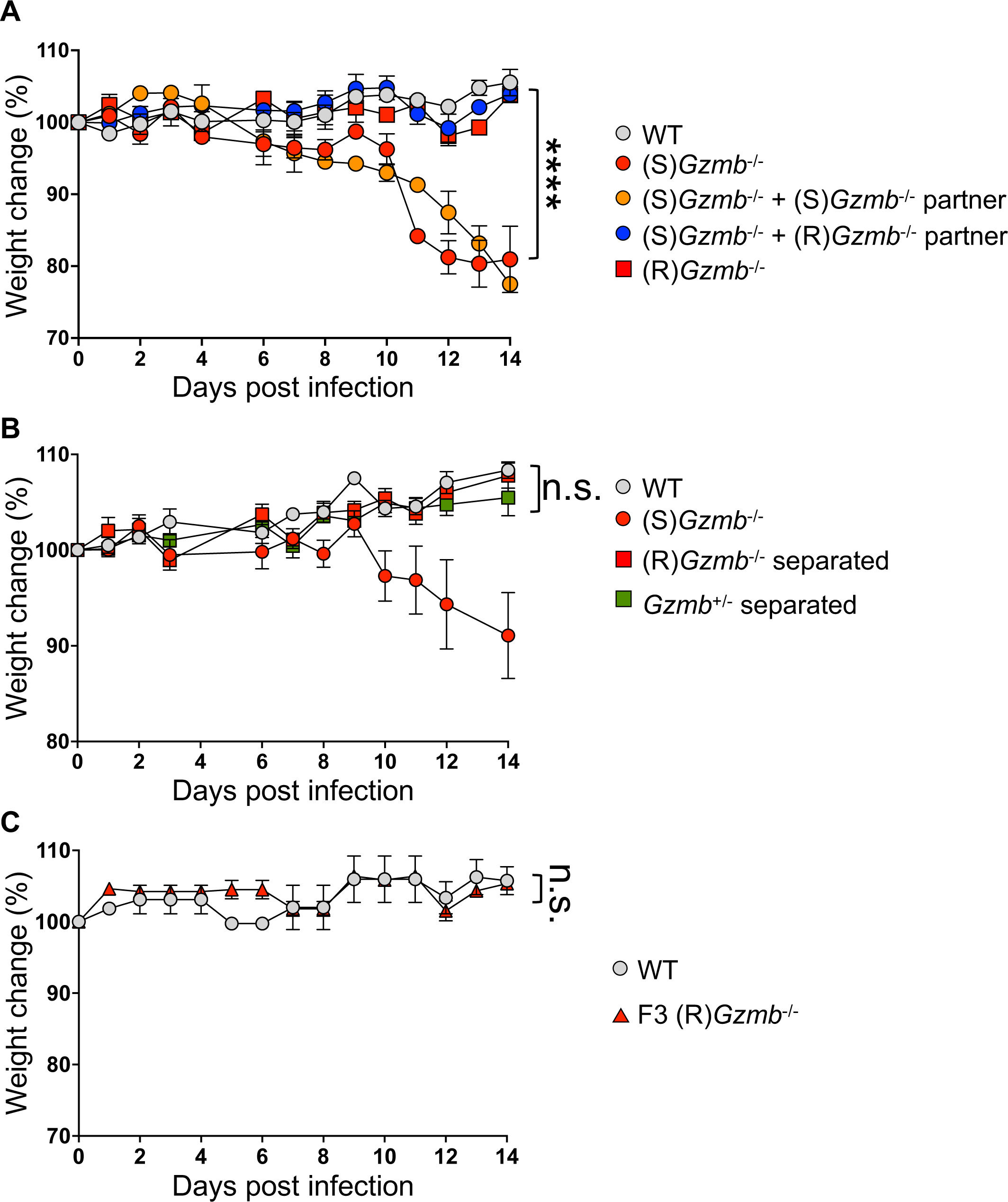
Resistance to *C. rodentium* is acquired by co-housing and maintained through time. (A) (S)*Gzmb*^-/-^ mice were co-housed with a single (S)*Gzmb*^-/-^ mouse from a different litter, or a single (R)*Gzmb*^-/-^ mouse starting at weaning age. Three to four weeks later, mice were infected with 5-10 × 10^8^ CFU of *C. rodentium* and were monitored for weight change. (B) Littermate (R)*Gzmb*^-/-^ and *Gzmb*^+/-^ mice were separated at weaning age and infected as in (A). (C) (R)*Gzmb*^-/-^ mice were bred among themselves. F3 mice were infected as in (A). n= 4-10. Data represent at least 3 independent experiments. For (A), ****P<0.001, Student’s t test for each time point after day 8. n.s.= not significant.

Susceptible *Gzmb*^-/-^ mice in **Figure 2A** were cohoused with at least one resistant mouse in the cage starting at weaning age, raising the possibility that *Gzmb*^-/-^ mice require the constant presence of a resistance-inducing microbiota to remain free of severe disease induced by *C. rodentium*. To test this possibility, we weaned and separated littermate (R)*Gzmb*^-/-^ and *Gzmb*^+/-^ mice based on sex and genotype. Mice were infected 7 to 8 weeks after birth. As shown in **Figure 2B**, *C. rodentium*-infected (R)*Gzmb*^-/-^ mice remained resistant to severe disease even when separated from *Gzmb*^+/-^ mice. To further investigate stability of the resistant phenotype, (R)*Gzmb*^-/-^ mice were bred among themselves, and their offspring were infected with *C. rodentium*. **Figure 2C** shows that F3 (R)*Gzmb*^-/-^ mice did not lose weight during the course of *C. rodentium* infection. Therefore, resistance to severe disease is maintained throughout the life span of the (R)*Gzmb*^-/-^ mice and can be transmitted to their offspring.

Because susceptibility to severe disease caused by *C. rodentium* infection was not transmissible by co-housing, we investigated whether this trait can be acquired by microbiota transplantation. For this purpose, WT, *Gzmb*^+/-^, and (S)*Gzmb*^-/-^ mice were treated with a broad spectrum antibiotic cocktail, followed by oral gavage with fecal matter derived from (S)*Gzmb*^-/-^ or (R)*Gzmb*^-/-^ mice. Mice were infected with *C. rodentium* 3 weeks post-fecal transplant. WT and *Gzmb*^+/-^ mice became susceptible to severe infection when transplanted with microbiota from (S)*Gzmb*^-/-^ mice (**Figures 3A-3D**). The resistance trait was maintained when transplanted with microbiota from (R)*Gzmb*^-/-^ mice, indicating that antibiotic treatment does not alter susceptibility to *C. rodentium* infection. Despite the differences in disease outcome, all mice displayed similar levels of *C. rodentium* colonization (**Figure 3E**). Overall, our results demonstrate that the intestinal microbiota is responsible for either resistance or susceptibility to severe disease caused by *C. rodentium*.

**Figure 3.**
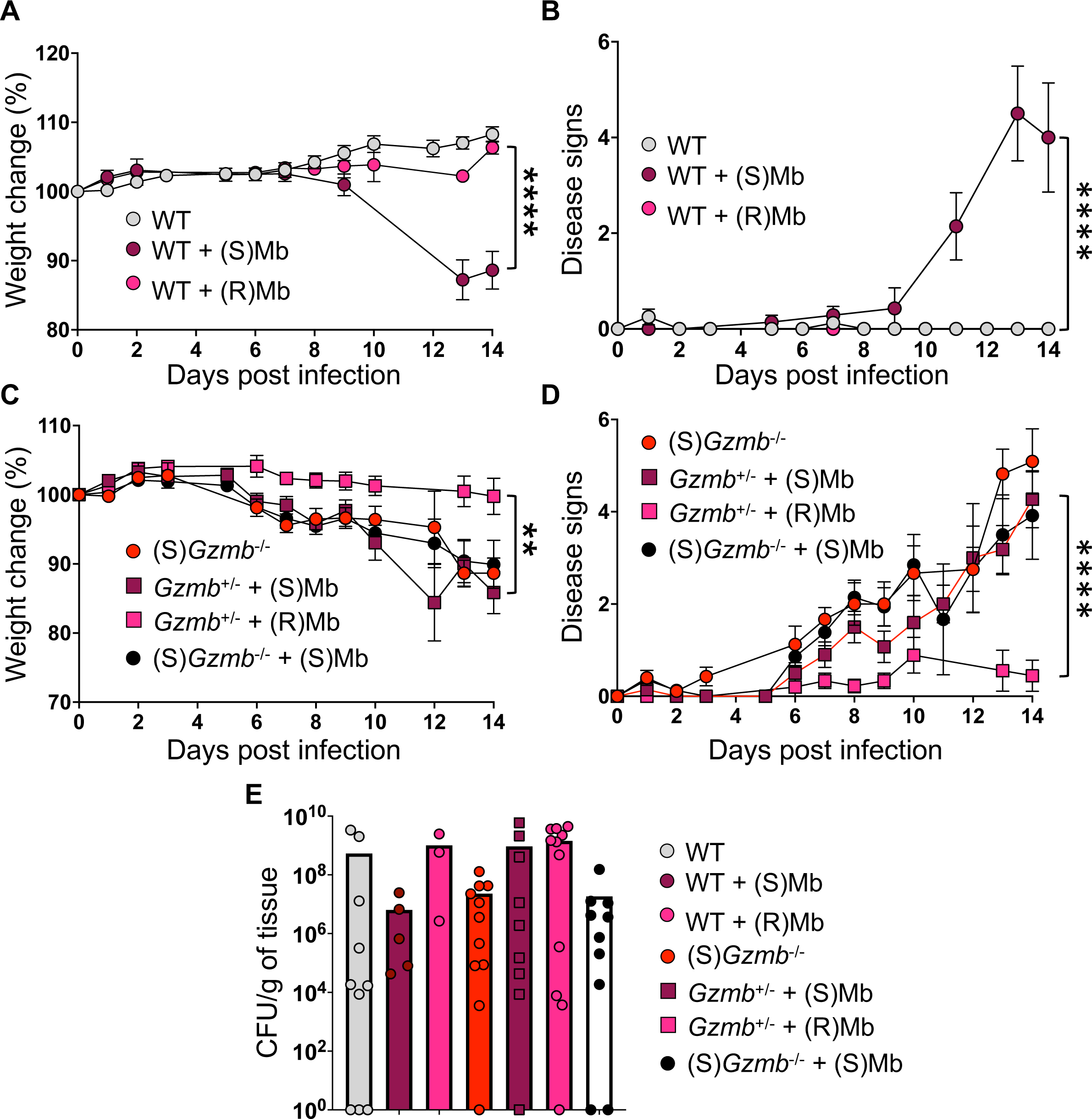
Resistance and susceptibility to severe *C. rodentium* infection are acquired by microbiota (Mb) transplant. The indicated mice were treated with a broad-spectrum antibiotic cocktail followed by fecal oral gavage from the indicated mice. Three weeks after microbiota transplant, mice were infected with 5-10 × 10^8^ CFU of *C. rodentium*. Mice were weighed (A and C) and monitored for disease signs (C and D). At the end point, *C. rodentium* colonization was determined. n= 5-12. Data is combined from at least 3 independent experiments. **P<0.01, Student’s t test for day 14; ****P<0.001, Student’s t test for day 14.

Analysis of the intestinal microbiota of (S)*Gzmb*^-/-^ and (R)*Gzmb*^-/-^ mice show similarities in the abundance of the most predominant bacterial orders (**Figure 4A**). The only apparent order with increased abundance in resistant mice, but almost absence in susceptible animals, was Turicibacterales (**Figure 4A and 4B**). This was confirmed by ANCOM (<u>an</u>alysis of <u>co</u>mposition of <u>m</u>icrobes) (**Figure 4C**).

**Figure 4.**
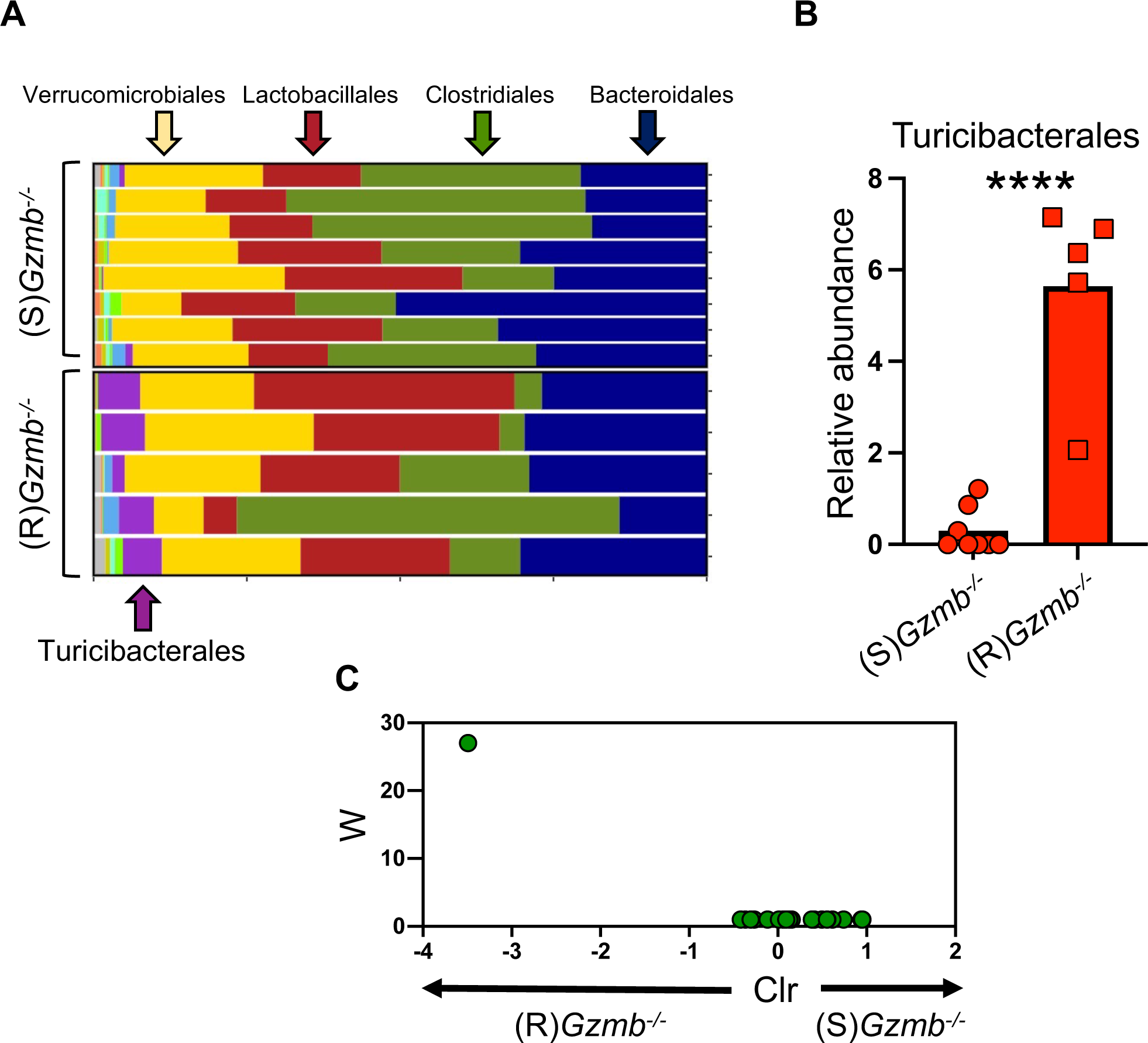
Turicibacterales are better represented in resistant mice. (A) 16S RNA gene sequence analysis from stool of the indicated naïve mice was analyzed for order abundance. (B) Relative abundance of the order Turicibacterales of the same samples as in (A). (C) ANCOM analysis. n= 5 -8. ****P<0.001, Student’s t test.

*Turicibacter sanguinis* is an anaerobic commensal, and one of the better studied species of Turicibacterales. However, the role of *T. sanguinis* as commensal is not known. Several isolates have been identified in human (such as, MOL361) and mice (H121, 1E2, TA25) (23, 24) *T. sanguinis* expresses a unique neurotransmitter sodium symporter-related protein with some homology to the mammalian serotonin transporter (20), which facilitates its identification by real-time PCR. To determine whether *T. sanguinis* is present in (R)*Gzmb*^-/-^ mice and absent in (S)*Gzmb*^-/-^ animals, we amplified the serotonin transporter gene from stool samples with isolate-specific primers, as previously reported (25). Mouse isolates H121 and 1E2 were detected in (R)*Gzmb*^-/-^ mice but not in susceptible animals (**Figure 5A**), while the human isolate MOL361 was absent in all mouse lines tested.

**Figure 5.**
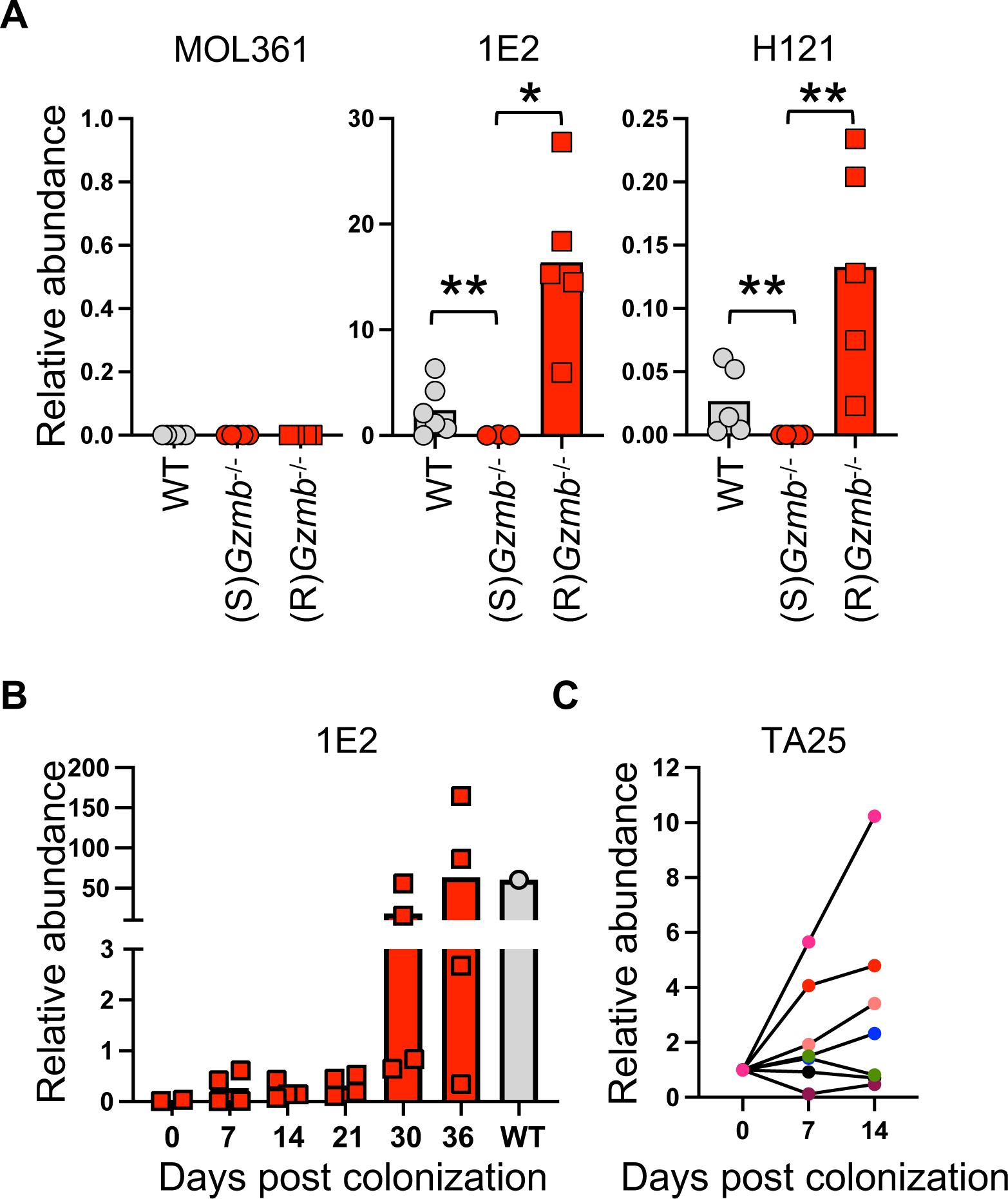
*T. sanguinis* is present in resistant *Gzmb*^-/-^ mice and is transferable. (A) Relative abundance of the *T. sanguinis* isolates in the feces of the indicated mice. Non-antibiotic treated (S)*Gzmb*^-/-^ mice were gavaged with stool content from resistant mice (B) or with cultured *T. sanguinis* TA25 (identified by the 1E2 primers) (C). Each symbol represents an individual mouse. n=4-6. *P<0.05; **p<0.01, Student’s t test.

Overall, these results indicate that one of the main differences in the intestinal microbial communities between (R)*Gzmb*^-/-^ and (S)*Gzmb*^-/-^ mice is the presence of *T. sanguinis* in the former and its absence in the latter. Moreover, these results suggest that this commensal may be responsible for conferring protection from severe disease caused by *C. rodentium*.

We have shown that resistance is transmitted by co-housing without the need of treating mice with antibiotics (**Figure 2A**), indicating that the bacterial communities conferring resistance can colonize without disrupting the native microbiota. To determine whether *T. sanguinis* can colonize intestines with established microbiota, non-antibiotic-treated (S)*Gzmb*^-/-^ mice were orally gavaged with fecal content from WT mice. *T. sanguinis* colonization was monitored in the stools at different time points after inoculation. *T. sanguinis* 1E2 was detected in most stool samples after 21 days post colonization (**Figure 5B**). Colonization of (S)*Gzmb*^-/-^ mice was evident at 7 days when mice were gavaged with *in vitro* grown cultures of *T. sanguinis* TA25, which can also be identified by the 1E2 primers (**Figure 5C**). These results demonstrate *T. sanguinis* colonizes undisrupted intestinal microbiota, and that gavage of *in vitro* grown *T. sanguinis* is a more efficient method of colonization by this commensal.

We next investigated whether colonization with *T. sanguinis* prevents severe disease in (S)*Gzmb*^-/-^ mice subsequently infected with *C. rodentium*. For this purpose, we colonized non-antibiotic-treated (S)*Gzmb*^-/-^ mice with cultured *T. sanguinis* TA25, and three weeks later, mice were infected with *C. rodentium*. Although (S)*Gzmb*^-/-^ mice colonized with *T. sanguinis* lost slightly more weight compared to WT mice, they were better protected than (S)*Gzmb*^-/-^ mice (**Figure 6A and 6B**). Despite less weight loss and decreased signs of disease, colon pathology and *C. rodentium* colonization of (S)*Gzmb*^-/-^ mice treated with *T. sanguinis* were similar to non-colonized (S)*Gzmb*^-/-^ mice (**Figure 6C and 6D**). *C. rodentium* localization within the intestinal epithelium was more prominent along the length of the crypts in (S)*Gzmb*^-/-^ mice, but not if they were colonized with *T. sanguinis* prior to infection (**Figure 7**). Whether this latter observation is responsible for the increased weight loss and disease severity in (S)*Gzmb*^-/-^ mice is currently unknown.

**Figure 6.**
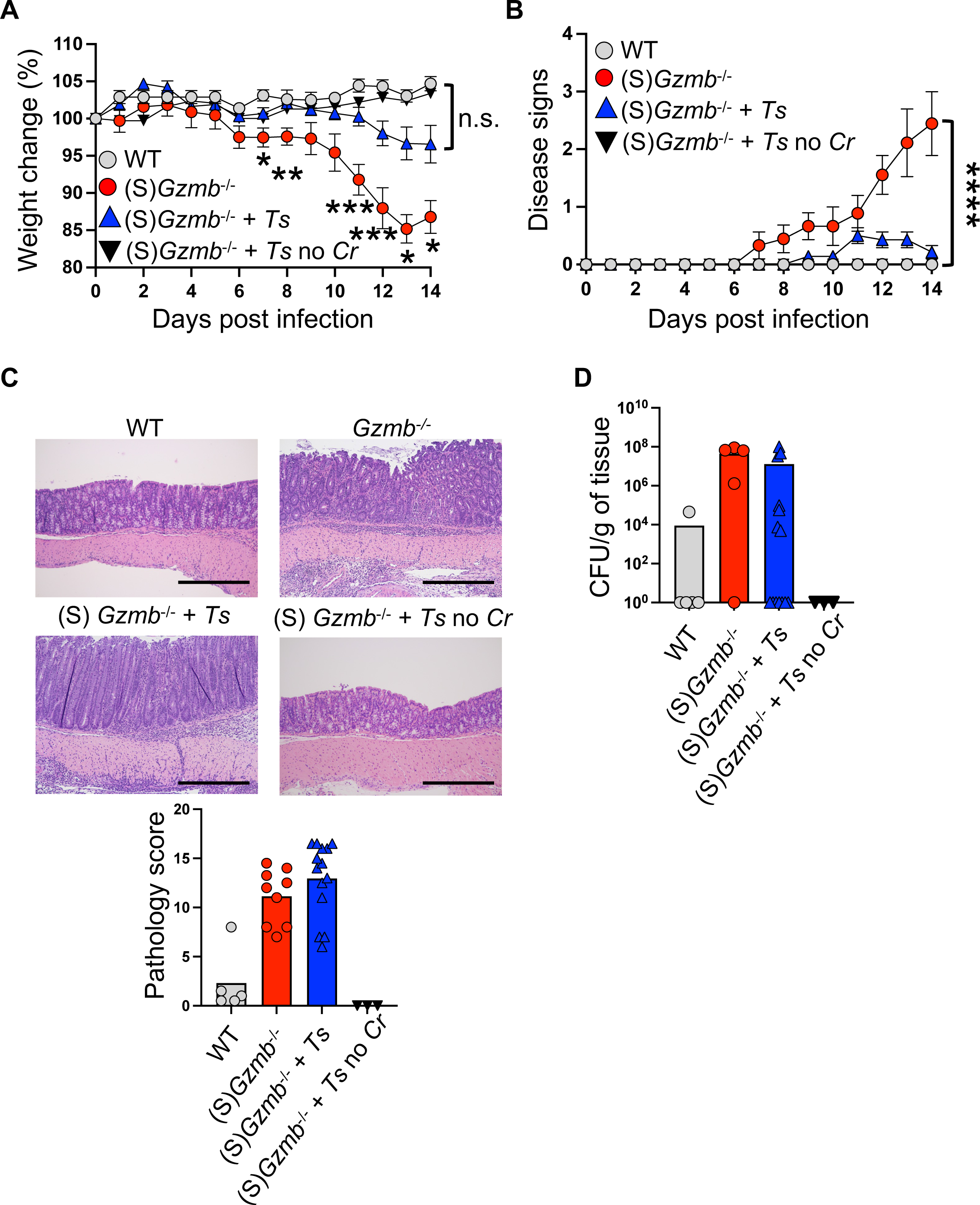
*T. sanguinis* confers partial protection from severe *C. rodentium* infection. Mice were colonized with 1-10 × 10^6^ CFU of *T. sanguinis* isolate TA25 3 times (d0, d3, and d7). Approximately 14 days later, mice were infected with 5-10 × 10^8^ CFU of *C. rodentium* and monitored for weight (A) and signs of disease (B). At the end point, colon pathology (C) and *C. rodentium* colonization (D) were determined. *Ts* = *T. sanguinis*; *Cr* = *C. rodentium*. Bar = 250mm. n=5-14. In bar graphs, each symbol represents and individual mouse. *P<0.05; **P<0.01; ***P<0.005; ****P<0.001; Student’s t test.

**Figure 7.**
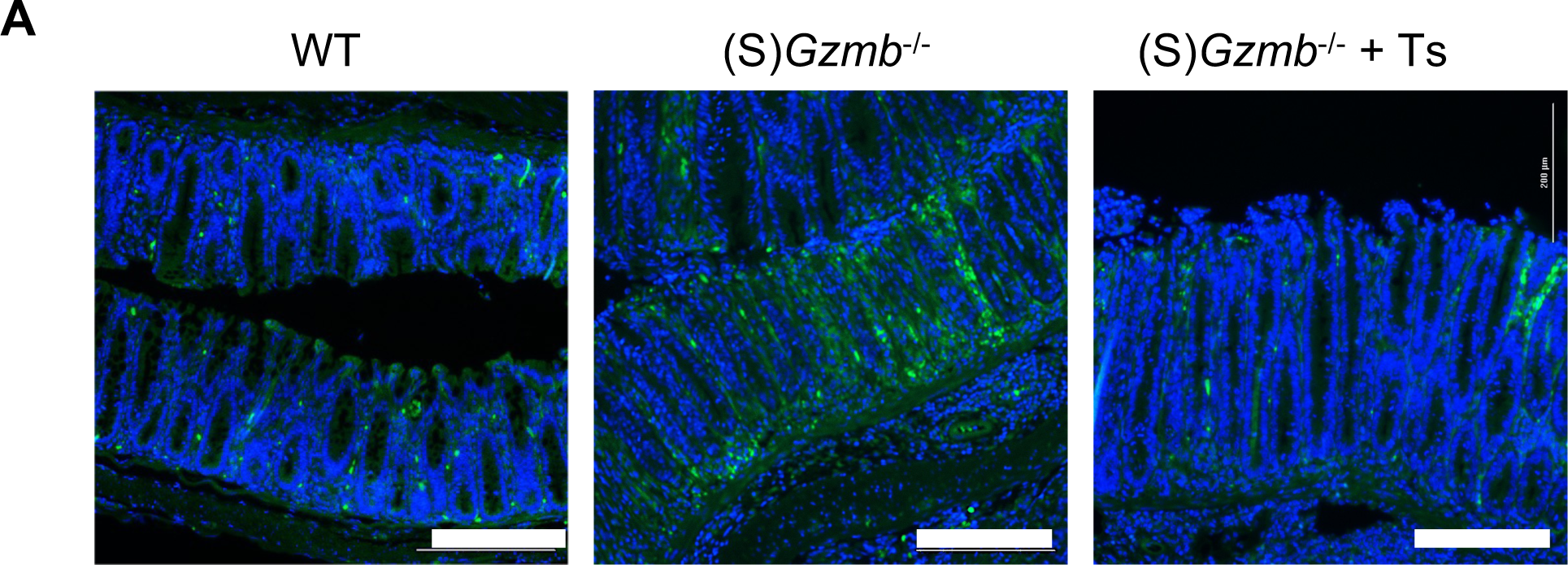
*C. rodentium* colonization in the indicated mice. Colon tissue from GFP-*C. rodentium* infected mice was fixed with 4% paraformaldehyde, paraffine embedded, and stained for DAPI. Samples were visualized using a Cytation C10 Confocal Imaging Reader and Gen 5+ software (Agilent BioTek). These samples are representative mice from Figure 6. Bar = 250mm.

Overall, the results presented in this section indicate that *T. sanguinis* colonization provides a protective effect from severe *C. rodentium* infection.

## DISCUSSION

Interventions for infectious diseases mostly concentrate on pathogen eradication. In the case of bacterial infections, antibiotics represent the first therapeutical tool to eliminate dangerous microorganisms. However, the use of antibiotics, though powerful, comes with many significant side effects including disruption of commensal communities. It has been known for a long time that alterations in the intestinal microbiota predisposes the host to colonization with pathogenic organisms (26). Therefore, a potential therapeutic intervention, especially for severe bacterial infections, is fecal microbiota transplantation. This technique aims at introducing potential beneficial communities that may hinder colonization of pathogenic microorganisms, or the expansion of pathobionts (27). Although mostly safe (28), one of the major risks involving fecal microbiota transplantation is the introduction of pathogens, which have caused severe disease and in some cases, death of the recipient individuals (29).

A better therapeutic approach might entail the introduction of specific commensal species with the capacity to restrain or control severe bacterial infections. Therefore, identification of commensals with these qualities represents a critical step towards the discovery of novel interventions. In this report, we present data indicating that *Turicibacter* represents a potential commensal involved in protection from severe intestinal infection. We show that *Gzmb*^-/-^ mice lacking *T. sanguinis* lose significant weight, develop diarrhea, display scruffiness, present increased colon pathology, and in some instances, some of these mice succumb to infection. This phenotype does not represent an artifact of the granzyme B-deficient mouse line, because susceptibility can be transferred to WT C57BL/6 mice by transplanting total fecal microbiota from mice lacking *T. sanguinis*. Thus, even genetically resistant mice can become susceptible to severe *C. rodentium* infection by altering their intestinal microbiota.

It is remarkable that *T. sanguinis* colonization can be achieved without treating the recipient mice with antibiotics. This indicates that niches for *T. sanguinis* are available in undisturbed intestinal microbiota, amplifying its potential as a therapeutic tool for severe intestinal infections.

In studies of young children with diarrhea or acute gastroenteritis, increased relative abundance of *Turicibacter* in stools was associated with healthy controls (30, 31). Although these reports do not establish causation between the presence of *Turicibacter* and children with no diarrhea, they suggest a potential association between *Turicibacter* and decreased susceptibility to diarrheal diseases. Therefore, our results indicating protection to severe *C. rodentium* infection in the presence of *T. sanguinis* establish a causation with important implications for diarrheal diseases in humans.

In our previous publication we reported that CD4^+^ T cells in granzyme B-deficient mice differentiate into highly pathogenic IL-17-producing cells, which promote higher disease severity in the T cell adoptive transfer model of colitis and during *C. rodentium* infection (22). Here we show that while *T. sanguinis*-colonized (S)*Gzmb*^-/-^ mice are resistant to severe *C. rodentium* infection, they still present mild levels of disease and colon pathology similar to that of non-colonized (S)*Gzmb*^-/-^ mice (**Figure 6A and 6C**). These results suggest that in granzyme B-deficient mice, both improperly differentiated CD4^+^ T cells and lack of *T. sanguinis* promote development of severe disease during *C. rodentium* infection. However, it cannot be ruled out that other commensals or their metabolites confer full protection from severe disease. We believe this may be the case because littermate *Gzmb*^-/-^ mice have reduced colon pathology similar to WT mice (**Figure 1C**), which contrasts with the increased colon pathology observed in infected *T. sanguinis*-colonized *Gzmb*^-/-^ mice (**Figure 6C**).

How *T. sanguinis* promotes protection from severe *C. rodentium* infection is not known. Here we show that although mice colonized with *T. sanguinis* present similar total *C. rodentium* colonization levels and colon pathology to non-colonized mice, severe disease is prevented. These results suggest that *T. sanguinis* does not control *C. rodentium* colonization, but instead may decrease its pathogenicity or skew the host’s mucosal immune system towards a non-detrimental response. We are currently investigating these possibilities.

Why do granzyme B-deficient mice lack *T. sanguinis*? Granzyme B is mostly known for its role in cell-mediated cytotoxicity (21), but it has been postulated that this enzyme also possesses extracellular roles (32). Indeed, unpublished data from our group shows that granzyme B can be detected in the lumen of the intestines. Thus, it is possible that lack of granzyme B renders an intestinal environment where *Turicibacter* may not easily thrive. However, our multigenerational studies in which *Gzmb*^-/-^ mice remain resistant to severe *C. rodentium* infection argue against the above possibility. The most likely scenario is that the original line of *Gzmb*^-/-^ mice lost *T. sanguinis* at some point. Because this line was maintained isolated, it never acquired this commensal. Studying the microbiota requires well-controlled experiments, including using littermates in the case of investigating host mutations. However, as this report shows, littermate controls may hide potential and interesting microbiota discrepancies.

In summary, we present evidence indicating that the presence of *T. sanguinis* in the intestinal microbiota results in increased protection from susceptibility to severe *C. rodentium* infection. Future work will focus on the mechanism(s) by which this commensal confers protection, and thus, will open new potential therapeutic options for the treatment of intestinal infections.

## MATERIALS AND METHODS

### Mice

C57BL/6 mice were originally purchased from The Jackson Laboratory (000664) and have been maintained and acclimated in our colony for several years. Granzyme B-deficient (*Gzmb*^-/-^) mice were kindly provided by Dr. Xuefang Cao. These mice were kept as isolated lines, unless otherwise specified. Littermate mice were generated by crossing *Gzmb*^-/-^ with WT mice to generate *Gzmb*^+/-^ mice, which were bred among themselves to generate *Gzmb*^+/+^, *Gzmb*^+/-^, and *Gzmb*^-/-^ littermates. For convenience, we bred littermate *Gzmb*^+/-^ and *Gzmb*^-/-^ mice, to generate mice used in some experiments. Littermates derived from female *Gzmb*^-/-^ and male *Gzmb*^+/-^ mice were weaned and caged based on sex but not genotype, unless otherwise indicated. Littermate mice remained together until the end points. For some experiments, littermate mice were separated at weaning age based on sex and genotype and remained separated until the end of experimentation. Some littermate *Gzmb*^-/-^ mice were bred among themselves to obtain F3. All mice were between 5 to 10 weeks of age at the time of experimentation, with *C. rodentium* infections occurring at 7-9 weeks of age. Male and female mice were used for all experiments. Mice were maintained in accordance with the Institutional Animal Care and Use Committee at Vanderbilt University Medical Center.

### Co-housing

At the time of weaning (3 to 4 weeks after birth), mice were separated by sex and a partner mouse of the same sex and age was added to the cage, remaining co-housed until the end of the experiment.

### Citrobacter rodentium *infection*

Seven to nine-week old mice were infected with *C. rodentium* (strain DBS100, ATCC 51459) or GFP-*C. rodentium* (12), as previously described (33). Results were similar and reproducible with either strain. As an example, experiments for Figure 1 were performed with the ATTC line, while those in Figure 6 were performed with the GFP-*C. rodentium* strain. Briefly, mice were infected with 5-10 × 10^8^ CFU exponentially grown bacteria by oral gavage. Starting weight was determined prior to gavage. Infection was carried out by Y.L.L. or K.L.H. Mice were weighed daily for 14 days. At the same time, mice were monitored for the following signs of disease: irritated rectum, loose stool, rectal bleeding, diarrhea, hunched posture, scruffiness, and moribund. Each sign was scored as one point and added. Weighing and monitoring after infection was done blindly by D.O.-V. At the end of the experiment, distal colons were isolated to determine bacterial colonization in MacConkey media (represented as CFU/g of tissue) and pathology performed blindly by Dr. Piazuelo as previously described (34). Colon tissue from GFP-*C. rodentium* infected mice was fixed with 4% paraformaldehyde, paraffine embedded, and stained for DAPI. Samples were visualized using a Cytation C10 Confocal Imaging Reader and Gen 5+ software (Agilent BioTek).

### Turicibacter sanguinis *colonization*

*T. sanguinis* isolate TA25 was kindly provided by Dr. Kenya Honda and Jonathan Lynch. *T. sanguinis* was grown in suspension under anaerobic conditions in Schaedlers broth. Mice were gavaged three times at d0, d3, and d7 with 1-10×10^6^ CFU. TA25 can be identified by the primers against 1E2 isolate.

### *Real time PCR* for detection of Turicibacter sanguinis

DNA was isolated from fecal pellets using the DNeasy Power Soil Kit (Qiagen) following the manufacturer’s instructions. Detection of the serotonin transporter (20), for isolates MOL361, 1E2, and H121 was performed as previously indicated (25). MOL361, forward GGGTTTGCAGATGCGGG, reverse AATTTATCAACCACCGCTGTAATAAT; 1E2, forward GGGTTTGCTGATGCCGC, reverse CCCAAATTTATCAACTACTGCTGTAATAAT; H121, forward GGTTTAGCTGATGCTGGAATTAG, reverse TCCAAATTTATCAACAAATGCTGTAATAAT

### Microbiota depletion and fecal transplant

Mice were treated with a cocktail of antibiotics as previously described (35, 36), with some modifications. Briefly, 5-week old mice were orally gavaged on day 0 and 1 with 200μl/mouse of the following antibiotic cocktail: ampicillin (2mg/ml), neomycin (2mg/ml), metronidazole (2mg/ml), vancomycin (1mg/ml) dissolved in sterile water. From day 0 to day 6, mice were provided a similar cocktail with half the concentration for each antibiotic in the drinking water. Tabletop sugar (50mg/ml) was added to the cocktail to improve antibiotic intake. Water containing fresh antibiotic cocktail was replenished at day 2 and 4 and was replaced with regular water at day 6. At day 8 and 10, 200 μl/mouse of fecal matter solution was gavaged as follows: At least 2 fecal pellets from donor mice were homogenized in 1ml of sterile PBS. Fecal solution was cleared by centrifugation at 8000g for 3 minutes. Supernatant was filtered through a 70μm mesh. At day 28, mice were infected with *C. rodentium* as indicated above.

### *16SrRNA* gene sequencing

At least 2 fecal pellets were collected per mouse and were stored at -80°C. DNA was isolated using the DNeasy Power Soil Kit (Qiagen, Germantown, MD, USA) following the manufacturer’s instructions. Extracted DNA concentration was measured using Nanodrop 2000 (Thermo Scientific, Waltham, MA, USA). The V3 and V4 hypervariable regions of the 16S rRNA gene were sequenced using 2 × 300 paired-end sequencing on the Illumina Miseq sequencing platform (Illumina) at Integrated Microbiome Resource, Dalhousie University, Halifax, NS, Canada. The QIIME2 pipeline (version 3.6.11) was used to process and filter demultiplexed sequence reads. OTUs were clustered using Deblur (2020.8.0) prior to alignment using QIIME2. OTU taxonomy was determined using a naive Bayesean classifier trained toward the GreenGenes 99% reference database (13_8). The OTU table was rarified to an even depth of references per sample prior to generation of taxonomy barplots using the ggplot2 and plotly packages in R (4.2.2). Analysis of Composition of Microbes (ANCOM) was performed using QIIME2. Analysis of Turicibacter relative abundance was calculated using Student’s t test in Graphpad Prism.

## ACKNOWLEDGEMENTS

This work was supported by NIH grants R01DK111671 (D.O.-V.) and R01DK128200 (K.T.W.); Vanderbilt Training in Cellular, Biochemical, and Molecular Sciences Training Program, T32GM008554-25 (K.G.M); National Science Foundation Graduate Research Fellowship under Grant No. 1937963 (K.G.M.); and the Digestive Disease Research Center at Vanderbilt University Medical Center, NIH grant P30DK058404 (M.B.P., K.T.W., and D.O.-V.). We acknowledge the Translational Pathology Shared Resource supported by NCI/NIH cancer center support grant 5P30CA68485-19 and the Mouse Metabolic Phenotyping Center Grant 2U24DK059637-16. We also acknowledge VA Merit Review grant I01CX002171 (K.T. W.) and Senior Research Award 703003 from the Crohn’s & Colitis Foundation (K.T. W.).

## References

1. Kotloff KL, Nataro JP, Blackwelder WC, Nasrin D, Farag TH, Panchalingam S, Wu Y, Sow SO, Sur D, Breiman RF, Faruque AS, Zaidi AK, Saha D, Alonso PL, Tamboura B, Sanogo D, Onwuchekwa U, Manna B, Ramamurthy T, Kanungo S, Ochieng JB, Omore R, Oundo JO, Hossain A, Das SK, Ahmed S, Qureshi S, Quadri F, Adegbola RA, Antonio M, Hossain MJ, Akinsola A, Mandomando I, Nhampossa T, Acacio S, Biswas K, O’Reilly CE, Mintz ED, Berkeley LY, Muhsen K, Sommerfelt H, Robins-Browne RM, Levine MM. 2013. Burden and aetiology of diarrhoeal disease in infants and young children in developing countries (the Global Enteric Multicenter Study, GEMS): a prospective, case-control study. Lancet 382:209–22.

2. Liu L, Johnson HL, Cousens S, Perin J, Scott S, Lawn JE, Rudan I, Campbell H, Cibulskis R, Li M, Mathers C, Black RE, Child Health Epidemiology Reference Group of WHO, Unicef. 2012. Global, regional, and national causes of child mortality: an updated systematic analysis for 2010 with time trends since 2000. Lancet 379:2151-61.

3. Croxen MA, Law RJ, Scholz R, Keeney KM, Wlodarska M, Finlay BB. 2013. Recent advances in understanding enteric pathogenic Escherichia coli. Clin Microbiol Rev 26:822–80.

4. Mundy R, MacDonald TT, Dougan G, Frankel G, Wiles S. 2005. Citrobacter rodentium of mice and man. Cell Microbiol 7:1697–706.

5. Ledwaba SE, Costa DVS, Bolick DT, Giallourou N, Medeiros P, Swann JR, Traore AN, Potgieter N, Nataro JP, Guerrant RL. 2020. Enteropathogenic Escherichia coli Infection Induces Diarrhea, Intestinal Damage, Metabolic Alterations, and Increased Intestinal Permeability in a Murine Model. Front Cell Infect Microbiol 10:595266.

6. Silberger DJ, Zindl CL, Weaver CT. 2017. Citrobacter rodentium: a model enteropathogen for understanding the interplay of innate and adaptive components of type 3 immunity. Mucosal Immunol 10:1108–1117.

7. Kim YG, Kamada N, Shaw MH, Warner N, Chen GY, Franchi L, Nunez G. 2011. The Nod2 sensor promotes intestinal pathogen eradication via the chemokine CCL2-dependent recruitment of inflammatory monocytes. Immunity 34:769–80.

8. Zheng Y, Valdez PA, Danilenko DM, Hu Y, Sa SM, Gong Q, Abbas AR, Modrusan Z, Ghilardi N, de Sauvage FJ, Ouyang W. 2008. Interleukin-22 mediates early host defense against attaching and effacing bacterial pathogens. Nat Med 14:282–9.

9. Dann SM, Spehlmann ME, Hammond DC, Iimura M, Hase K, Choi LJ, Hanson E, Eckmann L. 2008. IL-6-dependent mucosal protection prevents establishment of a microbial niche for attaching/effacing lesion-forming enteric bacterial pathogens. J Immunol 180:6816–26.

10. Mullineaux-Sanders C, Sanchez-Garrido J, Hopkins EGD, Shenoy AR, Barry R, Frankel G. 2019. Citrobacter rodentium-host-microbiota interactions: immunity, bioenergetics and metabolism. Nat Rev Microbiol 17:701–715.

11. Satoh-Takayama N, Vosshenrich CA, Lesjean-Pottier S, Sawa S, Lochner M, Rattis F, Mention JJ, Thiam K, Cerf-Bensussan N, Mandelboim O, Eberl G, Di Santo JP. 2008. Microbial flora drives interleukin 22 production in intestinal NKp46+ cells that provide innate mucosal immune defense. Immunity 29:958–70.

12. Bergstrom KS, Kissoon-Singh V, Gibson DL, Ma C, Montero M, Sham HP, Ryz N, Huang T, Velcich A, Finlay BB, Chadee K, Vallance BA. 2010. Muc2 protects against lethal infectious colitis by disassociating pathogenic and commensal bacteria from the colonic mucosa. PLoS Pathog 6:e1000902.

13. Feng Y, Wang Y, Zhu B, Gao GF, Guo Y, Hu Y. 2021. Metagenome-assembled genomes and gene catalog from the chicken gut microbiome aid in deciphering antibiotic resistomes. Commun Biol 4:1305.

14. Huang S, Ji S, Yan H, Hao Y, Zhang J, Wang Y, Cao Z, Li S. 2020. The day-to-day stability of the ruminal and fecal microbiota in lactating dairy cows. Microbiologyopen 9:e990.

15. Kim CY, Lee M, Yang S, Kim K, Yong D, Kim HR, Lee I. 2021. Human reference gut microbiome catalog including newly assembled genomes from under-represented Asian metagenomes. Genome Med 13:134.

16. Maki JJ, Looft T. 2022. Turicibacter bilis sp. nov., a novel bacterium isolated from the chicken eggshell and swine ileum. Int J Syst Evol Microbiol 72.

17. Yano JM, Yu K, Donaldson GP, Shastri GG, Ann P, Ma L, Nagler CR, Ismagilov RF, Mazmanian SK, Hsiao EY. 2015. Indigenous bacteria from the gut microbiota regulate host serotonin biosynthesis. Cell 161:264–76.

18. Mo J, Gao L, Zhang N, Xie J, Li D, Shan T, Fan L. 2021. Structural and quantitative alterations of gut microbiota in experimental small bowel obstruction. PLoS One 16:e0255651.

19. Browne HP, Forster SC, Anonye BO, Kumar N, Neville BA, Stares MD, Goulding D, Lawley TD. 2016. Culturing of ‘unculturable’ human microbiota reveals novel taxa and extensive sporulation. Nature 533:543–546.

20. Fung TC, Vuong HE, Luna CDG, Pronovost GN, Aleksandrova AA, Riley NG, Vavilina A, McGinn J, Rendon T, Forrest LR, Hsiao EY. 2019. Intestinal serotonin and fluoxetine exposure modulate bacterial colonization in the gut. Nat Microbiol 4:2064–2073.

21. Anthony DA, Andrews DM, Watt SV, Trapani JA, Smyth MJ. 2010. Functional dissection of the granzyme family: cell death and inflammation. Immunol Rev 235:73–92.

22. Hoek KL, Greer MJ, McClanahan KG, Nazmi A, Piazuelo MB, Singh K, Wilson KT, Olivares-Villagomez D. 2021. Granzyme B prevents aberrant IL-17 production and intestinal pathogenicity in CD4(+) T cells. Mucosal Immunol doi:10.1038/s41385-021-00427-1.

23. Auchtung TA, Holder ME, Gesell JR, Ajami NJ, Duarte RT, Itoh K, Caspi RR, Petrosino JF, Horai R, Zarate-Blades CR. 2016. Complete Genome Sequence of Turicibacter sp. Strain H121, Isolated from the Feces of a Contaminated Germ-Free Mouse. Genome Announc 4.

24. Bosshard PP, Zbinden R, Altwegg M. 2002. Turicibacter sanguinis gen. nov., sp. nov., a novel anaerobic, Gram-positive bacterium. Int J Syst Evol Microbiol 52:1263–1266.

25. Lynch JBG, E.L.; Choy, K.; Faull, K.F.; Jewell, T.; Arellano, A.; Liang, J.; Yu, K.B.;, Paramo, J.; and Hsiao, E.Y. 2022. Turicibacter modifies host bile acids and lipids in a strain-specific manner. BioRxiv doi:https://doi.org/10.1101/2022.06.27.497673.

26. Bohnhoff M, Drake BL, Miller CP. 1954. Effect of streptomycin on susceptibility of intestinal tract to experimental Salmonella infection. Proc Soc Exp Biol Med 86:132–7.

27. Merrick B, Allen L, Masirah MZN, Forbes B, Shawcross DL, Goldenberg SD. 2020. Regulation, risk and safety of Faecal Microbiota Transplant. Infect Prev Pract 2:100069.

28. Baxter M, Colville A. 2016. Adverse events in faecal microbiota transplant: a review of the literature. J Hosp Infect 92:117–27.

29. DeFilipp Z, Bloom PP, Torres Soto M, Mansour MK, Sater MRA, Huntley MH, Turbett S, Chung RT, Chen YB, Hohmann EL. 2019. Drug-Resistant E. coli Bacteremia Transmitted by Fecal Microbiota Transplant. N Engl J Med 381:2043–2050.

30. Pop M, Walker AW, Paulson J, Lindsay B, Antonio M, Hossain MA, Oundo J, Tamboura B, Mai V, Astrovskaya I, Corrada Bravo H, Rance R, Stares M, Levine MM, Panchalingam S, Kotloff K, Ikumapayi UN, Ebruke C, Adeyemi M, Ahmed D, Ahmed F, Alam MT, Amin R, Siddiqui S, Ochieng JB, Ouma E, Juma J, Mailu E, Omore R, Morris JG, Breiman RF, Saha D, Parkhill J, Nataro JP, Stine OC. 2014. Diarrhea in young children from low-income countries leads to large-scale alterations in intestinal microbiota composition. Genome Biol 15:R76.

31. Quaye EK, Adjei RL, Isawumi A, Allen DJ, Caporaso JG, Quaye O. 2023. Altered Faecal Microbiota Composition and Structure of Ghanaian Children with Acute Gastroenteritis. Int J Mol Sci 24.

32. Afonina IS, Cullen SP, Martin SJ. 2010. Cytotoxic and non-cytotoxic roles of the CTL/NK protease granzyme B. Immunol Rev 235:105–16.

33. Olivares-Villagomez D, Algood HM, Singh K, Parekh VV, Ryan KE, Piazuelo MB, Wilson KT, Van Kaer L. 2011. Intestinal epithelial cells modulate CD4 T cell responses via the thymus leukemia antigen. J Immunol 187:4051–60.

34. Singh K, Gobert AP, Coburn LA, Barry DP, Allaman M, Asim M, Luis PB, Schneider C, Milne GL, Boone HH, Shilts MH, Washington MK, Das SR, Piazuelo MB, Wilson KT. 2019. Dietary Arginine Regulates Severity of Experimental Colitis and Affects the Colonic Microbiome. Front Cell Infect Microbiol 9:66.

35. Zhang Q, Lu Q, Luo Y. 2020. Fecal Microbiota Transplantation and Detection of Prevalence of IgA-Coated Bacteria in the Gut. J Vis Exp doi:10.3791/60772.

36. Bokoliya SC, Dorsett Y, Panier H, Zhou Y. 2021. Procedures for Fecal Microbiota Transplantation in Murine Microbiome Studies. Front Cell Infect Microbiol 11:711055.

